## Supplemental Fig A for "Comment on Pescott & Jitlal 2020: Failure to account for measurement error undermines their conclusion of a weak impact of nitrogen deposition on plant species richness"

Supplementary Figure A: Estimated mean of the random spatial field fitted to the data using the SPDE approach in INLA.

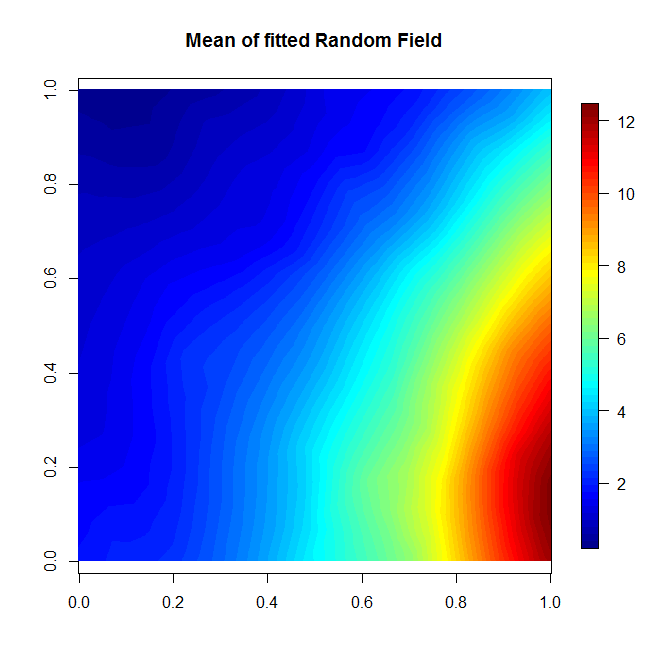
