## Supplementary material for "Comment on Pescott & Jitlal 2020: Failure to account for measurement error undermines their conclusion of a weak impact of nitrogen deposition on plant species richness": R code for simulation study

####################################################################################
####################################################################################
##
##  Author: Pete Henrys , April 2020
##
##  Code for simulation study to show effects of 
##  measurement error in spatial attribution models
##
##  Smart et al., 2020. Comment on Pescott & Jitlal 2020: Failure to account for 
##  measurement error undermines their conclusion of a weak impact of nitrogen 
##  deposition on plant species richness
##
##
####################################################################################
####################################################################################


## Load the required libraries
library(fields)
library(INLA)

#create empty matrices to store parameter estimates and DIC values
sim.out=sim.out2=matrix(NA,nrow=100,ncol=7)
diccomp=matrix(NA,nrow=100,ncol=2)


#loop over 100 iterations of the simulation
for(ki in 1:100){

#####################
##    Simulate Data
#####################

#define a grid on the unit square on which to simulate data 
grd <- expand.grid(x=seq(0,1,len=100),y=seq(0,1,len=100))

#define a deterministic hypothetical spatial covariate - denoted SpCov in text
grd$sp.cov <- exp(2*grd$x - 1.2*(grd$y^2))

#based on this covariate, define a hypothetical response variable as a linear function of the spatial covariate eplus random noise
grd$resp <- 10 + 2*grd$sp.cov + rnorm(length(grd[,1]),0,0.5)

#plot the relationship between the covariate and response for visualisation (shown in Figure 1b)
plot(grd$sp.cov,grd$resp)

#plot the pattern of the spatial covariate (shown in Figure 1a)
colr=tim.colors(10)
plot(grd$x,grd$y,pch=15,cex=0.7,col=colr[ceiling(grd$sp.cov)])
 
#create a coarse grid on unit square to reflect spatial aggregation of covariate  
grd.big <- expand.grid(x=seq(0,1,len=5),y=seq(0,1,len=5))

#specifiy distance function that will be used to align covariates and grid cells 
distm <- function(x0,x1,y0,y1){
	x=sqrt((x0-x1)^2 + (y0-y1)^2)
	return(which.min(x))
}

##match coarse grid cells with original fine scale grid cells
mn = integer(length(grd[,1]))
for(j in 1:length(grd[,1])){
	mn[j] <- distm(grd$x[j],grd.big$x,grd$y[j],grd.big$y)
}

#calculate the mean value of the covariate for each aggregated cell and add some random noise
mn.cov <- tapply(grd$sp.cov,mn,mean) + rnorm(length(unique(mn)),0,0.5)

#specify coarse spatial covariate (denoted SpCovErr in the main text) by aligning original cells (ie in grd) all with the aggregated cells (ie in grd.big) within which they sit 
grd$sp.cov.crse <- mn.cov[mn]

#randomly sample 100 grid cells with which to run analyses
idx <- sample(1:length(grd[,1]),100)
dat <- grd[idx,]

#plot the relationship between the response variable and the aggregated covariate with error (shown in Figure 1c)
plot(dat$sp.cov.crse,dat$resp)


#####################
##    Modelling
#####################


# define mesh upon which to estimate the spatial random field using the SPDE approach
mesh <- inla.mesh.2d(loc=rbind(dat[,1:2]),max.edge=c(0.1,0.2),cutoff=0.05)

#plot the mesh to ensure that the node density looks reasonable
plot(mesh)

#specify spde model - here priors are assigned to the matern hyper parameters 
spde <- inla.spde2.pcmatern(mesh,prior.range=c(0.5,0.5),prior.sigma=c(5,0.5))

#create the A matrix to enable conversion from observed locations to mesh
KFD_A <- inla.spde.make.A(mesh=mesh,loc=as.matrix(dat[,1:2]))

#make the inla stack
stk_KFD <- inla.stack(data=list(y=dat$resp),
						effects=list(data.frame(sp.cov.crse=dat$sp.cov.crse,intercept=1), 
						Bnodes=1:spde$n.spde),
						A=list(1,KFD_A),
						tag="est.KFD")	

#specify full formula for model including covariate and spatial random field
formula = y ~  intercept + sp.cov.crse +
						f(Bnodes, model = spde) - 1  

#specify formula for linear model excluding spatial field
formlin = y ~  intercept + sp.cov.crse - 1  						
						
#fit the full model in INLA ensuring that the DIC is returned 
result <- inla(formula,
				data=inla.stack.data(stk_KFD),
					control.predictor=list(link=1,A=inla.stack.A(stk_KFD),compute=TRUE),
					control.compute = list(dic = TRUE),
					verbose=FALSE
								)
								
#fit the linear model in INLA ensuring that the DIC is returned 
reslin <- inla(formlin,
				data=inla.stack.data(stk_KFD),
					control.predictor=list(link=1,A=inla.stack.A(stk_KFD),compute=TRUE),
					control.compute = list(dic = TRUE),
					verbose=FALSE
								)								
								
# Extract the estimated fixed effects from this model 
res <- result$summary.fixed
res2 <- reslin$summary.fixed

# Plot the estimate mean of the random field 
#first by creating a projection between the mesh and a grid over whcih to predict the field
projg<-inla.mesh.projector(mesh,ylim=range(grd$y,na.rm=T),xlim=range(grd$x,na.rm=T),dims=c(500,1000))
#then estimating the mean of the field at these locations across the grid
xmean1 <- inla.mesh.project(projg, result$summary.random$Bnodes$mean)
#and finally, producing an image plot of the surface
image.plot(seq(min(grd$x,na.rm=T),max(grd$x,na.rm=T),len=500),seq(min(grd$y,na.rm=T),max(grd$y,na.rm=T),len=1000),xmean1, col.regions=tim.colors(),xlab='', ylab='', scales=list(draw=FALSE),main="Mean of fitted Random Field",asp=1)


#store the estimated coefficient corresponding to the spatial covariate
sim.out[ki,] <- as.numeric(res[2,])
sim.out2[ki,] <- as.numeric(res2[2,])

#store the DIC of both models
diccomp[ki,] <- c(result$dic$dic,reslin$dic$dic)

print(ki)

}
